# Imaging and tracking mRNA in live mammalian cells via fluorogenic photoaffinity labeling

**DOI:** 10.1101/2020.02.10.942482

**Authors:** Tewoderos M. Ayele, Travis Loya, Arielle N. Valdez-Sinon, Gary J. Bassell, Jennifer M. Heemstra

## Abstract

Cellular RNA labeling using light-up aptamers that bind to and activate fluorogenic molecules has gained interest in recent years as an alternative to protein-based RNA labeling approaches. Aptamer-based systems are genetically encodable and cover the entire visible spectrum. However, the relatively weak nature of the non-covalent aptamer-fluorogen interaction limits the utility of these systems in that multiple copies of the aptamer are often required, and in most cases the aptamer must be expressed on a second scaffold such as a transfer RNA. We propose that these limitations can be averted through covalent RNA labeling, and here we describe a photoaffinity approach in which the aptamer ligand is functionalized with a photoactivatable reactive group such that irradiation with UV light results in covalent attachment to the RNA of interest. In addition to the robustness of the covalent linkage, this approach benefits from the ability to temporally control RNA labeling. To demonstrate this method, we incorporated a photoaffinity linker onto malachite green and fused the malachite green aptamer to a specific mRNA reporter of interest. We observed markedly improved sensitivity for fixed cell imaging of mRNA using this approach compared to *in situ* hybridization. Additionally, we demonstrate visualization of RNA dynamics in live cells using an mRNA having only a single copy of the aptamer, minimizing perturbation of the structure and localization. Our initial biological application utilizes the photoaffinity labeling approach to monitor RNA stress granule dynamics and we envision future application of this method for a wide range of investigations into the cellular localization, dynamics, and protein binding properties of cellular RNAs.

---

Trafficking of messenger RNA (mRNA) to subcellular compartments plays an essential role in RNA homeostasis and cellular function. This spatiotemporal control of mRNA localization is a common characteristic for a significant fraction of transcripts,^(1-3)^ and in recent years, fluorescent microscopy has dramatically increased our understanding of the heterogeneity of transcript regulation and the complex subcellular interactions of RNAs and proteins. However, this relies on the ability to fluorescently label cellular RNAs without significantly perturbing their structure or localization. The earliest approaches to fluorescently tagging cellular RNAs utilized probes capable of binding to the RNA of interest (ROI) through Watson-Crick-Franklin base pairing, including fluorescent *in situ* hybridization (FISH) and molecular beacons. While these methods yielded much of the current day knowledge on RNA localization, they generally require cell fixation, and thus cannot provide insight into trafficking and dynamics of cellular RNAs.^(4-6)^ Currently, the most ubiquitous approach for visualizing mRNA uses GFP-fused RNA binding proteins such as MS2, λ_N_, PCP, or Cas proteins.^(7-10)^ These fluorescently-tagged proteins recognize a specific sequence that is incorporated multiple times onto ROI. While this does enable visualization of RNAs in living cells, these methodologies suffer from the fact that the unbound fluorescent protein creates significant background signal. This necessitates functionalization of the ROI with multiple copies of the target RNA sequence, and the size of that sequence as well as the heavy load of the associated proteins (>1300 kDa) can alter the native localization and functional properties of the RNA.^(11, 12)^

In 2003, Tsien and coworkers proposed that the small-molecule recognition capabilities of RNA could potentially be used for RNA labeling, and they reported an RNA aptamer that binds to the fluorogenic dye malachite green (MG). Over the past decade, a number of other RNA aptamers binding to fluorogenic dyes have been reported, including the Spinach^(13, 14)^, Broccoli^(15)^, and Mango^(16)^ aptamers. These aptamers have been fused to RNAs and expressed in cells to enable RNA visualization. However, these approaches remain predominantly used in bacterial cells and typically require fusion of multiple copies of the aptamer to enable fluorescence imaging. Recently, Yang and coworkers reported the Peppers aptamer system, which has a high signal-to-background ratio and was used to image genomic loci in mammalian cells. While the stability and cellular brightness of the Peppers aptamer was a significant improvement compared to the previously reported aptamer-based systems, this approach still suffers from inherent limitations as a result of its non-covalent nature. For example, the reversibility of the fluorescent probe interaction with the ROI makes Peppers and other non-covalent systems unusable for applications that require media exchange, as labeling does not withstand washout steps. This inherent limitation of non-covalent approaches also makes them unsuitable for time-resolved investigations such as pulse-chase experiments.

We hypothesized that the limitations of the current RNA labeling approaches could be overcome through covalent attachment of the fluorescent molecule to the target RNA. Compared to all of the existing methods, which rely on non-covalent binding, covalent attachment would provide increased robustness to maximize signal-to-background and would allow the labeling to withstand media exchange or washing steps. Additionally, we envisioned that using a photoactivatable reactive group would provide temporal control over the labeling process, which is not possible using any of the existing methods. To achieve this goal, we utilized the malachite green aptamer (MGA) first reported by Tsien and coworkers.^(17)^ Similar to the Peppers aptamer, MGA binds to the MG fluorogen and induces significant red-shifted fluorescence enhancement.^(18)^ The excitation and emission maxima for the MG fluorogen are also located in the far-red region of the UV spectrum, averting the inherent challenges associated with cellular auto-fluorescence and making this aptamer-ligand pair exceptionally well-suited for live cell imaging.

To covalently label the target RNA, we envisioned that the aptamer could be fused to the ROI, expressed in cells, and the cells incubated with the MG ligand having a photoactivatable handle. UV-irradiation would convert the non-covalent binding interaction into a covalent linkage, resulting in robust and temporally-controlled labeling of the ROI (Fig. 1a). To create the photo-reactive fluorogen, we appended a diazirine linker to the dye (Fig. 1b) and termed this new MG derivative malachite green diazirine-2 (MGD2). We previously used a similar approach to design a derivative of MG termed MGD for photoaffinity labeling of proteins in live cells;^(19)^ however, this strategy has yet to be applied to RNA labeling. Upon UV-A irradiation at 365 nm, the diazirine linker is activated and produces a carbene moiety that readily reacts with nearby C-H and heteroatom-H bonds. UV-C (254 nm) irradiation has been used for cross-linking RNA-protein interactions by taking advantage of the photo-responsiveness of natural amino acids such as Cys, Lys, Phe, Trp, and Tyr.^(20, 21)^ However, the longer wavelength of 365 nm used to activate MGD2 ensures that our design does not result in unwanted cross-linkage of RNA with cellular proteins.

**Fig. 1.**
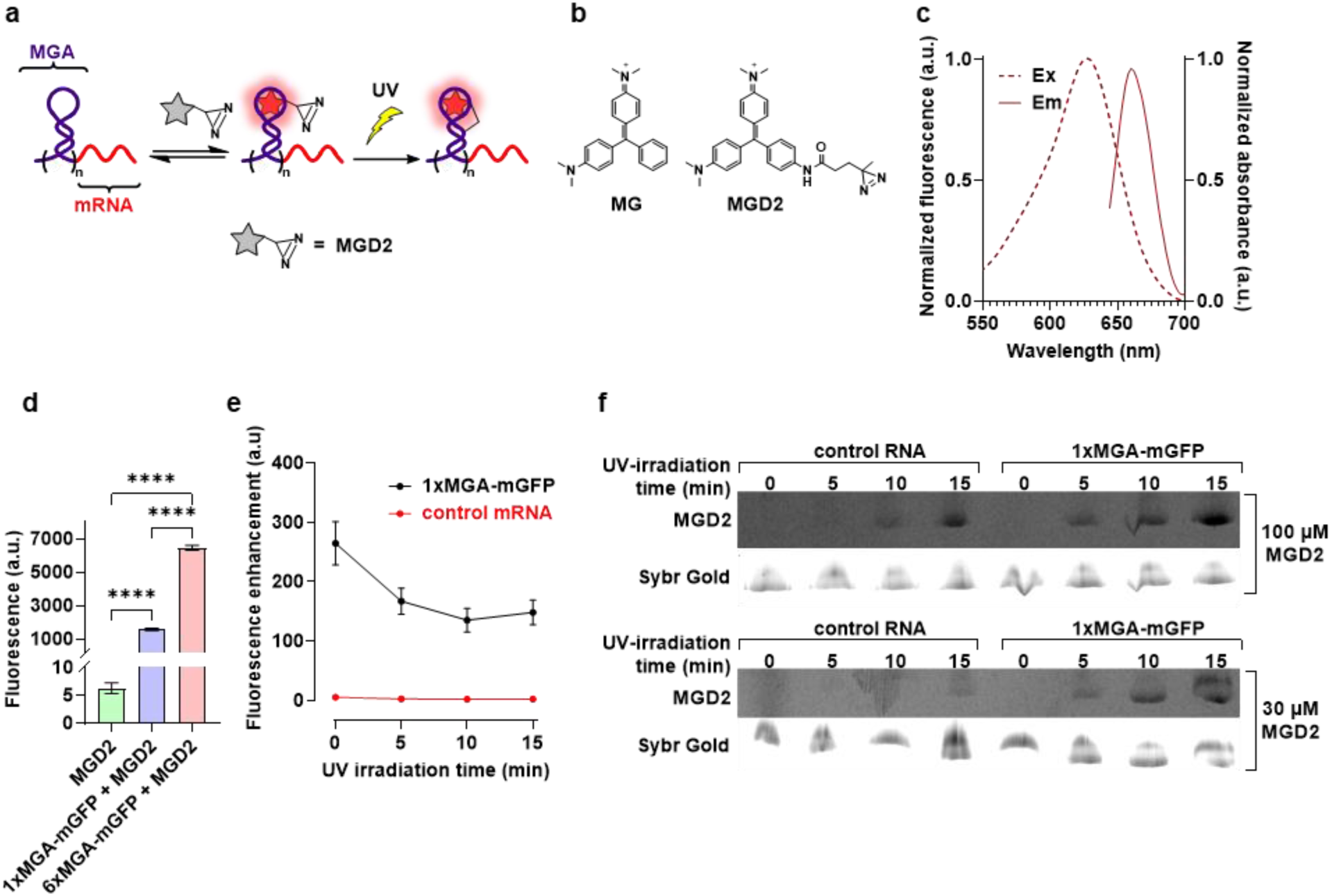
Characterization of MGA-functionalized mRNA in the presence of MGD2. **a**, Schematic representation of fluorogenic photoaffinity labeling of MGA-functionalized mRNA. The MGA-functionalized mRNA binds to the fluorogenic dye and induces fluorescence enhancement. UV-irradiation results in covalent attachment of the dye to the ROI. **b**, Structures of MG and MGD2. The canonical MG molecule was functionalized with a diazirine linker to enable photoaffinity labeling of MGA. **c**, Emission (solid) and excitation (dashed) spectra of MGD2 bound to 1xMGA-mGFP. **d**, Fluorescence of MGD2, 1xMGA-mGFP, and 6xMGA-mGFP in 1xPBS. Statistical comparison was performed using one-way ANOVA. (****P<0.0001). Bars and error bars represent the mean and standard deviation from n = 4 independent samples. **e**, Time-dependent fluorescence enhancement of 1xMGA-mRNA and control mRNA upon irradiation at 360 nm in 1x PBS. Points and error bars represent the mean and standard deviation from n = 3 independent samples. **f**, Denaturing PAGE gel analysis of UV-dependent covalent labeling of 1x MGA-mGFP and control GFP mRNAs with MGD2.

Using our photoaffinity approach, we demonstrate that an mRNA of interest can be labeled and imaged in both fixed and live cells using a single 57 nt fusion. This is significantly smaller than the fusions required in other aptamer-based methods, minimizing perturbation of RNA structure and localization. We demonstrate that covalent labeling enables RNA visualization under conditions where the non-covalent system fails, and we utilize our approach to monitor the dynamics of RNAs in stress granules. Together, this research introduces the first covalent method for cellular RNA labeling and provides an effective and easy-to-use tool for the RNA community. The added robustness and temporal control achieved using this approach is anticipated to significantly advance RNA imaging capabilities, providing important new insights into the role of RNA trafficking in biological processes such as development and disease.

## Results

### *In vitro* characterization

We first utilized *in vitro* studies to investigate the reactivity and selectivity of the aptamer when functionalized to an mRNA. This was accomplished by transcribing the acGFP mRNA appended with one or six copies of the aptamer sequence at the 5’ UTR, which will be referred to as 1xMGA-mGFP and 6xMGA-mGFP, respectively. Both MGA-functionalized mRNAs displayed a well-defined absorbance and fluorescence profile with an excitation maximum at 625 nm and an emission maximum 660 nm in the presence of MGD2 (Fig. 1c). Prior to UV-irradiation, fluorescence measurements revealed a 251-fold fluorescence enhancement for MGD2 bound to 1xMGA-mGFP and >1000-fold fluorescence enhancement for MGD2 bound to 6xMGA-mGFP (Fig. 1d). We next investigated the effect of UV-irradiation on the fluorescence enhancement. For this experiment, we used 1xMGA-mGFP and a control mGFP mRNA that does not contain the aptamer sequence (Fig. 1e). We observed that up to 15 minutes of UV-irradiation did not result in any detectable fluorescence enhancement of MGD2 in the presence of the control mRNA. However, the fluorescence enhancement of 1xMGA-mGFP declined slightly over time and stabilized at approximately 140-fold after 15 minutes of UV irradiation. This decrease in enhancement was not entirely unexpected, as covalent attachment may limit the fluorogen to binding modes that have slightly less rotational restriction, but nonetheless we remained encouraged by these data as they indicated that a single copy of MGA could produce a detectable and stable signal-to-background ratio for cellular imaging experiments.

We also investigated the specificity of labeling using denaturing PAGE analysis. Both the control GFP mRNA lacking MGA (4 µM) and 1xMGA-GFP mRNA (4µM) were incubated with MGD2 and UV-irradiated for different lengths of time (Fig. 1f). We observed non-specific labeling of the control mRNA after 15 minutes of irradiation in the presence of 30 µM MGD2 and after 10 minutes of irradiation in the presence of 100 µM MGD2. These data were somewhat surprising given the lack of fluorescence enhancement observed for the control RNA in the previous experiment. However, this observation can be explained by considering that in the case of non-specific labeling, the energy from the absorbed light is dissipated through nonradiative rotational relaxation of the phenyl groups of MGD2.^(17, 22)^ However, the tight target-specific binding of the aptamer restricts the rotational relaxation of the dye and results in the enhanced fluorescence output. Thus, while our approach does result in some non-specific RNA labeling, we anticipated that this would not create problematic background signal during imaging experiments. We were further encouraged to this notion upon testing the selective reaction of MGD2 (30 µM) with 6xMGA-mGFP in the presence of cellular RNA (Supplementary Fig. 1). After 10 min of UV irradiation, we observed bands in the MG channel corresponding to the target RNA and slight impurities, but no bands from labeling of other cellular RNAs. The collective observations from these experiments served as guidelines for irradiation time and dye concentrations used in subsequent cellular labeling experiments.

### Fixed-cell imaging of RNA

Having demonstrated our photoaffinity labeling approach *in vitro*, we next turned to fixed cell experiments, as this would enable us to directly compare our method to FISH and validate labeling of the target RNA in cells. As a biological context for testing our labeling method, we chose stress granules. In response to stress conditions, cells form non-membrane-bound cytosolic and nuclear RNA-protein assemblies to stall the translation of mRNA until the cells are no longer under stress. While most mRNAs can be concentrated to stress granules, different mRNAs have vastly distinct localization efficiencies.^(5, 23, 24)^ Using FISH, Parker and coworkers showed that the CDK6 mRNA is highly enriched in stress granules of mammalian cells.^(5)^ Inspired by this finding, we used mCDK6 as a model system to fluorescently label and image the unique distribution pattern of the mRNA. Both the 6x and 1x MGA sequences were inserted at the 5’ UTR of the transcript, and the construct was placed under the control of the cytomegalovirus (CMV) promoter (Fig. 2a). Mouse neuroblastoma Neuro-2a cells were then transfected with these plasmids and incubated with 30 µM MGD2 for 20 min. The cells were then irradiated using 365 nm UV light to allow for covalent labeling of the aptamer. The media was replaced to washout excess dye, and cell stress conditions were induced by 45 min of arsenite exposure, a well characterized paradigm to induce formation of stress granules. Following fixation and immunofluorescence labeling of one stress granule marker protein (G3BP1), the cells were imaged using confocal microscopy.

**Fig. 2.**
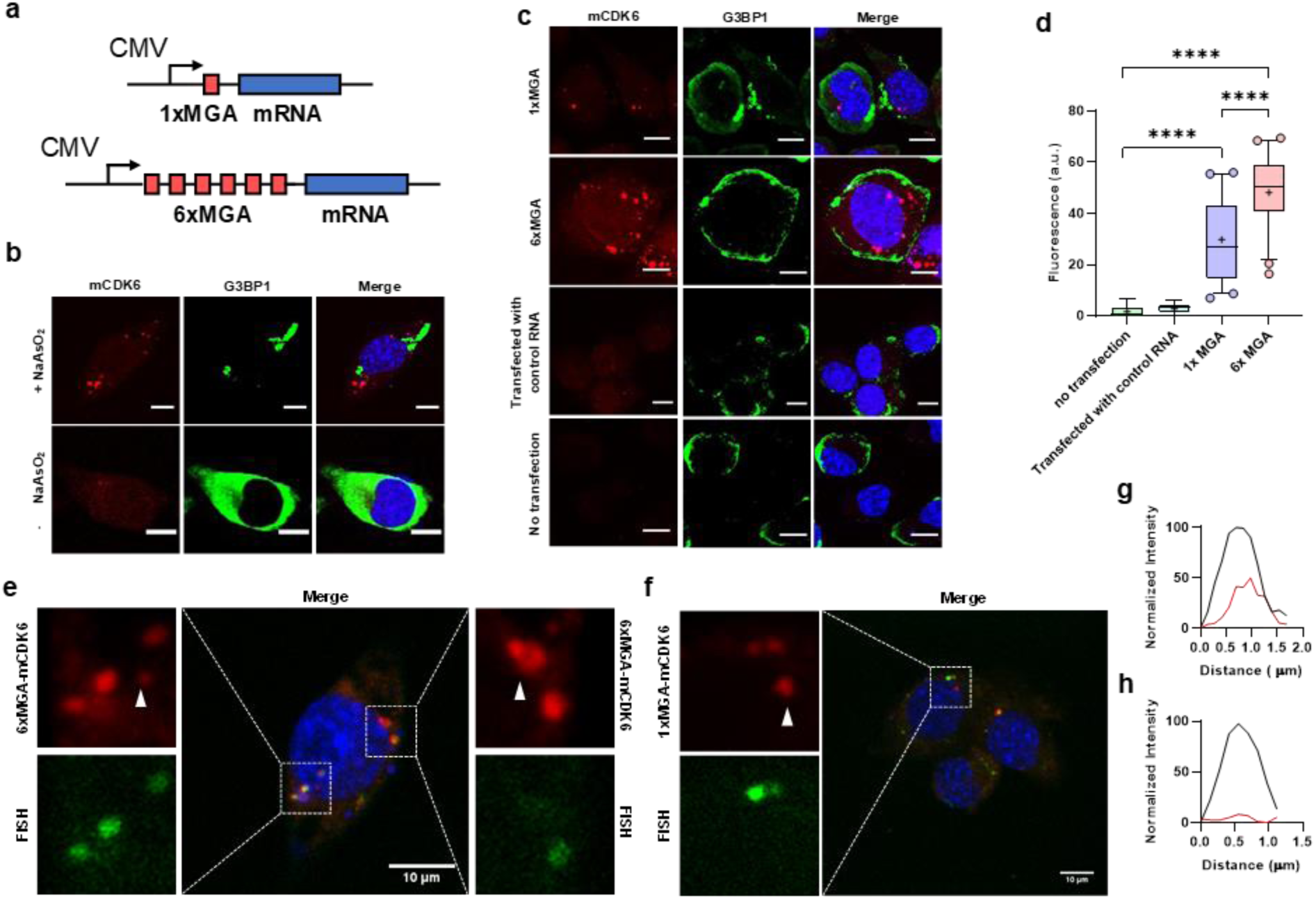
RNA labeling in fixed cells. **a**, Schematic representation of MGA-functionalized mRNA constructs. **b**, Representative confocal images of cellular granule formation in Neuro-2a cells with and without NaAsO2 stress. For this experiment, 6xMGA was used to label mCDK6 RNA. **c**, Representative confocal imaging of Neuro-2a cells transfected with 1xMGA-mCDK6, 6xMGA-mCDK6, control mCDK6, and cells that were not transfected. For figures 2b and c, G3BP1 protein immunolabeling was used to see the formation of granules. **d**, Fluorescence intensity of RNA foci in untransfected Neuro-2a cells or Neuro-2a cells expressing mCDK6 functionalized with 1xMGA or 6xMGA at the 5’UTR. (n = 6 foci for no transfection and transfection with control RNA, n = 50 foci for 1xMGA and 6xMGA). Statistical comparison was performed using one-way ANOVA. (****P<0.0001). Box plots show median, upper and lower quartiles, whiskers extending to 5^th^ and 95^th^ percentile, and mean represented by a cross sign. **e**, Confocal mCDK6 RNA labeling in Neuro-2a cells using 6xMGA and FISH. **f**, Confocal mCDK6 RNA labeling in Neuro-2a cells using 1xMGA and FISH. White arrow represents RNA granules detected by 6xMGA but not by FISH. Scale bars represent 10 µm. **g**, Representative normalized line-scan of colocalized FISH (red line) and MGA-mCDK6 (black line) labeling with MGD2. **h**, Representative normalized line-scan of RNA granules detected by MGA-functionalized mCDK6 (black line) but not with FISH (red line).

In Neuro-2a cells exposed to arsenite stress, we observed the formation of distinct cytoplasmic RNA granules with the 6xMGA-functionalized mCDK6 (6xMGA-mCDK6). In contrast, the signal from cells that were not exposed to arsenite was diffused uniformly throughout the cytoplasm, and no detectable stress granule enriched mCDK6 was observed (Fig. 2b). Encouraged by the ability to visualize RNA granules having six copies of the aptamer appended to the mRNA, we attempted to image cells that were transfected with 1xMGA-functionalized mCDK6 (1xMGA-mCDK6). After similar arsenite treatment, we observed that RNA granules could be detected with a single copy of the aptamer fused to the mRNA. Moreover, when arsenite-treated cells were not UV-irradiated, no detectable foci formation was observed, indicating that RNA visualization is dependent on covalent attachment of the probe to the aptamer. We were especially excited by this observation, as it validated our hypothesis that covalent RNA labeling would provide a more robust imaging method than the existing non-covalent approaches. To further validate the specificity of this system, we transfected cells with CDK6 lacking the MGA sequence and then incubated the cells with MGD2 and performed UV irradiation. These control cells did not show any labeling, indicating the absence of non-specific labeling of other cellular components (Fig. 2c). When comparing the stress granules detected with 1xMGA versus 6xMGA-functionalized mRNA, we did observe reduced fluorescence intensity in cytoplasmic mRNA granules (Fig 2c and d). However, the difference was only ∼2-fold compared to the 6-fold smaller fusion of the 1xMGA construct, and the ability to detect RNA localization using a single aptamer fusion allows for minimal alteration of the target mRNA. Together, these results demonstrate that we are able to fluorescently label cellular mRNAs in a sequence-specific manner and observe their localization to stress granules.

### Comparison of fixed cell imaging of MGA/MGD2 with FISH

To validate that the fluorescence signal observed was arising from labeling of the target CDK6 mRNA, we simultaneously incubated the Neuro-2a cells with FISH probes complementary to the CDK6 sequence but bearing a spectrally orthogonal fluorophore. This experiment also enabled us to directly compare the sensitivity of the MGA/MGD2 system to the commonly used FISH technique for RNA labeling in fixed cells. Following arsenite stress, MGD2 labeling, and cell fixation, we incubated the cells with our custom FISH probes. Merged image analysis of the 1xMGA- and 6xMGA-labeled mRNA with the FISH signal showed an overlap of the fluorescence generated from these two approaches (Fig. 2e-g). Interestingly, we observed some RNAs by our MGA/MGD2 system that were not identified by FISH (Fig. 2e, f, and h). This enhanced sensitivity was observed for both 1xMGA- and 6xMGA-functionalized mRNAs, indicating that our approach has a higher sensitivity for RNA detection than FISH. We reason this is because during FISH labeling, the fluorescent probe is hybridized after the ROI is localized to granules. While this approach can identify ROIs that are spatially accessible to the FISH probes, other proteins and RNAs found within the granules compromise the ability of the probes to hybridize with the ROI. In contrast to FISH, our approach ensures that the MGA-functionalized mRNA is labeled with the fluorescent reporter before it is localized to the granules. This important distinction in the timing of RNA labeling allows for the detection of mRNA that otherwise would be inaccessible for FISH labeling.

### Live cell imaging of mRNA

Having established the applicability and robustness of our approach for both *in vitro* and fixed cell RNA imaging, we next investigated whether the MGD2/MGA system would be suitable for live cell imaging of endogenous mRNA localization and dynamics. Moreover, we sought to determine whether we could simultaneously track the real-time localization properties of both RNA and proteins within a living cell. For this assay, we co-transfected Neuro-2a cells with expression plasmids for 1xMGA-mCDK6 and dual GFP-tagged G3BP1 protein. The expressed mRNA was labeled with MGD2 and photo-crosslinked. The cells were then imaged to record the spatiotemporal features of both mRNA and protein in stress granules. After arsenite exposure, real-time confocal microscopy imaging of the granules revealed the dynamic nature of both the mRNA and the protein (Fig 3a and Supplemental video 1). Monitoring the signal intensity, we observed the fluorescent signal generated from the MGD2 labeled granules remained consistent over >25 min of imaging. However, the GFP signal showed a sharp signal decrease after 17 min of imaging (Fig 3a and b). Time-lapse imaging of the RNA granules also showed a gradual phase separation and maturation of RNA granules (Fig 3c). These data highlight the ability of our approach to enable live-cell monitoring of RNA dynamics using only a single copy of the aptamer fusion and demonstrate the high photostability of the MG-aptamer pair.

**Fig. 3.**
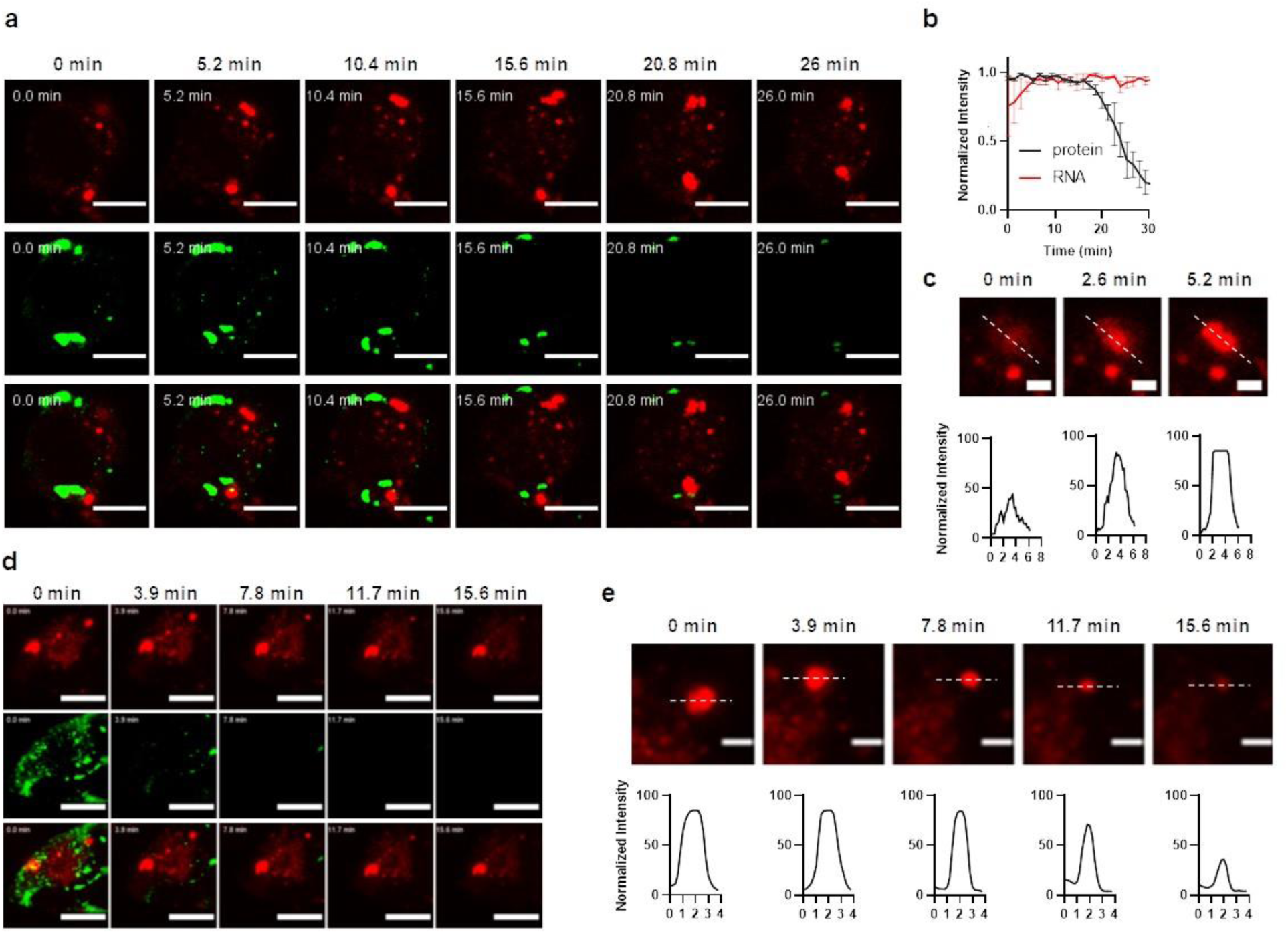
Live cell tracking of RNA and protein granules. **a**, Real-time confocal microscopy images showing dynamic localization of mCDK6 (top panel), G3BP1 protein (middle panel), and merged (bottom panel) in Neuro-2a cells with 5.2 min intervals. Scale bars represent 10 µm. **b**, Fluorescence signal comparison of MGD2-labeled mRNA vs GFP-labeled G3BP1 protein. Line graph represents the mean and error bars represent SEM from n = 3 granules. **c**, Image showing phase separation and RNA granule maturation. Top panel shows image slices taken every 2.8 min, and bottom panel is line-scan showing relative intensity of the dotted lines. Scale bar represents 2 µm **d**, Real-time confocal microscopy images showing granule deliquescence of mCDK6 (top panel), G3BP1 protein (middle panel), and merged (bottom panel) upon the addition of 5% 1,6-Hexanediol. Time slices were taken every 3.9 min. Scale bar represents 10 µm. **e**, Disappearance of RNA granule with 5% 1,6-Hexandiol treatment. Top panel shows confocal microcopy image of RNA granule and bottom panel shows relative intensity line-scan of the dotted lines. Scale bars represent 2 µm

After observing the motility and maturation of the granules, we questioned whether we could also observe their dissolution. In mammalian cells, stress granules disassemble in the presence of small aliphatic molecules that disrupt weak hydrophobic interactions.^(25)^ Therefore, we added 1,6-hexanediol, an aliphatic alcohol commonly used for disassembly of stress granules.^(25, 26)^ After arsenite-induced stress granule formation, we incubated the cells in 5% 1,6-hexanediol solution and observed the fluorescence signal of the labeled 1xMGA-mCDK6. In this experiment, signal from the protein granules disappeared after 7 min of incubation with 5% hexanediol. In contrast, most of the RNA granules exhibited a more sustained structural integrity, indicating that granules having high RNA content may be more resistant to dissolution (Fig 3d). In some RNA granules, however, hexanediol triggered an observable dissolution of these phase-separated compartments (Fig 3e and Supplemental video 2). This observation indicates that the strength of intermolecular forces of the RNA granule components is heterogeneous across different granules. Although we did not further investigate this property of RNA granules, the heterogeneity of the intermolecular forces of RNA granules is reported to be highly dependent on the composition of RNA, well-folded domains of proteins, and proteins having intrinsically disordered regions.^(27)^ Therefore, these data indicate that our RNA labeling approach is able to validate previously held assumptions of RNA properties as well as uncover lesser-known physical and biological characteristics of these biomolecules.

## Conclusion

Fluorescent labeling and imaging of RNA is key to understanding its roles in cellular function and disease processes. While a number of methods have been reported for labeling RNA, these all suffer from limitations related to either the requirement for cell fixation, the need for a large fusion added to the RNA, or lack of robustness due to the weak nature of non-covalent interactions. We sought to develop a broadly applicable strategy that can be implemented for both fixed cell and live cell imaging and that would enable robust labeling with only a single small RNA fusion and fluorophore tag. To achieve these goals, we recognized that all of the current methods rely on non-covalent binding interactions, and that covalent tethering of the fluorescent probe and the RNA of interest could provide increased signal-to-background ratio with a smaller RNA fusion. Specifically, we envisioned a photoaffinity labeling approach, as this would also provide temporal control over the RNA labeling process. We modified the malachite green fluorogen to incorporate a photo-reactive diazirine linker, which allowed for covalent labeling of its cognate aptamer upon irradiation with UV light. By placing this aptamer at the 5’ UTR of the mRNA, we showed target-specific fluorescence enhancement and labeling of the ROI. Fixed cell imaging of aptamer-functionalized mRNA showed formation of RNA stress granules in response to arsenite exposure. Compared to hybridization-based RNA labeling, we obtained enhanced sensitivity and lower background signal with our MGD2/MGA system. Furthermore, we showed that the dynamics of RNA granules containing a single aptamer-functionalized ROI can be tracked in live cells upon covalent attachment of the fluorogenic probe.

This novel strategy provides several advantages for RNA imaging applications. First, the far red-shifted fluorescence emission wavelength and the fluorogenic nature of the MGD2 dye allows for minimal background signal generated from cellular autofluorescence and unbound small molecules, respectively. Second, the temporally controlled covalent attachment of the fluorogen to its cognate aptamer enables the labeling to withstand washout steps and allows tracking of RNAs labeled during a specific time window, a feature which is necessary for pulse-chase studies and other experiments that require media exchange. Third, a single aptamer fusion of 57 nt was sufficient to image RNA granules in both live and fixed cells. This is significantly smaller than the fusions required in other RNA labeling approaches, which typically append numerous copies of the respective tag and fluorescent probe or proteins, producing a fusion that can add thousands of kDa. Finally, this strategy is anticipated to be highly generalizable to enable the development of additional aptamer-photoaffinity probe combinations for multiplexed and multicolor imaging. Together, the research reported here significantly advances RNA labeling technology and introduces a robust and reliable tool for use by the RNA community to study basic mechanisms that underlie localization and dynamics of diverse types of RNA granules and how these mechanism go awry in disease models and other important biological contexts.

## Methods

### *In vitro* fluorescence enhancement

A solution of 30 µM of MGD2 was mixed with 4 µM of the corresponding mRNA in 1x PBS (1.54 mM Potassium monobasic, 155.17 mM Sodium Chloride, and 2.71 mM Sodium Phosphate dibasic with pH = 7.2) (ThermoFisher) and incubated at room temperature for 20 min. For comparison, a solution of MGD2 without any RNA was also prepared in 1x PBS. The fluorescence of these solutions was measured using a BioTek Cytation 5 plate reader with ex = 620 ± 20 nm and em = 680 ± 20 nm. The average fluorescence value from replicate experiments was used to calculate fluorescence enhancement (*f*(x)) using Eq. 1, where F_MGA-mRNA_ is the fluorescence of the solution containing both MGA-fused mRNA and MGD2 and F_MGD2_ is the fluorescence of the MGD2 solution.

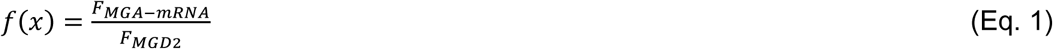

### Cell culture and transfection

Neuro-2a cells were used for all cellular experiments. These cells were cultured in DMEM (high glucose, Gibco) supplemented with 10% FBS at 37 °C in a humidified atmosphere of 5% CO_2_ and 95% air. The cells were split every two days or once they reached > 85% confluency. All cellular imaging was done on Cellview cell culture slides (Greiner Bio-One cat. No. 543079). All transfections were done using the Lipofectamine 3000 (Invitrogen) transfection reagent following the manufacturer recommended protocol with minor modifications: a solution of 5 µl of Opti-MEM media (Gibco) and 0.3 µl of Lipofectamine 3000 was mixed with a solution of 5 µL of Opti-MEM, 0.3 µg of DNA, and 0.8 µL of P3000. This solution was incubated at room temperature for 20 min. The media from the cell culture wells was removed and the DNA lipofectamine mixture was added directly to each chamber containing 60-80% confluent cells. Immediately after, 90 µl of 37 °C warmed media was added to each well for a total of 100 µL of solution. The cells were then placed back into the cell culture incubator for 12 h before conducting further experimentation.

### RNA FISH and MGA/MGD2 co-imaging

RNA FISH probes were designed against mCDK6 by using the Stellaris^®^ RNA FISH Probe Designer (Biosearch Technologies, Inc) available online at www.biosearchtech.com/stellarisdesigner (Version 4.2). The synthesis of these probes was done in-house using a solid-phase oligonucleotide synthesizer (MerMade 12). The probes were then labeled with Fluorescein using the Label IT^®^ nucleic acid labeling reagent (Mirus) using the manufacturer recommended protocol. Cells were incubated with 20 µM MGD2 in 37 °C prewarmed media for 20 min and irradiated with UV light at 365 nm for 10 min. The cells were then incubated in 0.5 mM sodium arsenite for 45 min at 37 °C. Cells wells were fixed with 4% paraformaldehyde (Biotium) for 15 min at room temperature and permeabilized with 0.1 % Triton X-100 (Sigma) for 1h. FISH probes were hybridized to the target mRNA for 12 h at 37 °C using Stellaris^®^ hybridization buffer containing 10% formamide. Cells were washed with 200 µL Wash Buffer A (Stellaris^®^) for 30 min in the dark followed by a wash with 200 µL Wash Buffer B for 5 min. Cells were then imaged in Vectashield^®^ antifade mounting media with DAPI (Vector Laboratories).

### Immunofluorescence and MGA/MGD2 co-imaging

Immunofluorescence and MGA/MGD2 imaging on fixed Neuro-2α cells were performed following transfection, MGA/MGD2 labeling, and cell fixation protocol described in the **RNA FISH and MGA/MGD2 co-imaging** methods section above. After fixing the MGD2 labeled cells with 4% formaldehyde, the cells were permeabilized for 1 h using blocking buffer containing: 5% rabbit serum (Millipore), 0.1% bovine serum albumin (Millipore), and 0.1% Triton (Sigma) in 1x PBS. The blocking buffer was then removed and replaced with 1:200 (v/v) diluted primary rabbit anti-G3BP1 antibody (Cell Signaling Technology^®^ # 17798). The primary antibody was diluted in dilution buffer containing: 1% bovine serum albumin (Millipore), and 0.1 % Triton X-100 (Sigma) in 1x PBS. After 12 h incubation with the primary antibody buffer solution, the buffer was removed and cells were washed for three times, 5 min each, with 1x PBS. The cells were then incubated in a solution of secondary goat anti-rabbit IgG H&L (Alexa Fluor^®^ 488) antibody (abcam #150077) for 1 h. The secondary antibody solution was prepared by diluting the secondary antibody to 1:200 (v/v) in the same dilution buffer as above. The cells were then washed three times for 5 min each with 1x PBS and imaged in Vectashield^®^ antifade mounting media with DAPI (Vector Laboratories).

### Confocal microscopy

Live and fixed cell images were taken on Cellview cell culture slides (Greiner Bio-One cat. No. 543079) using a Leica SP8 confocal laser scanning microscope equipped with an HC Plan Fluotar x10/0.15 air objective, an HC PL APO CS2 x20/0.75 air objective, an HC PL APO CS2 x63/1.4 oil objective, and two HYD GaAsP detectors. 405 nm Argon laser excitation was used to image DAPI; 488 nm Argon laser excitation was used to image Alexa488 labeled secondary antibody and FAM labeled FISH probes; 633 nm Helium-Neon laser was used to image MGD2.

Cellular images were obtained by taking Z-stack images with the instrument optimized step size and enough steps to cover the depth of each cell. Gain for each channel was optimized to minimize oversaturation while obtaining a clear fluorescent foci signal above background.

### Live cell imaging

Cells were transfected with Phage Ubic G3BP1-GFP-GFP which was gifted from Jeffery Chao (Addgene plasmid # 119950; http://n2t.net/addgene:119950; RRID:Addgene_119950). Twelve hours after transfection, the cells were incubated in 30 µM MGD2 in media for 20 min followed by 10 min UV-irradiation. The media was then removed and replaced with 37 °C prewarmed media. The live cell images were acquired using the same microscope and settings outlined above with the following modifications: the 6-well cell culture slides were placed in an environmental chamber to control humidity and temperature during imaging, and images were taken using a 60x oil-immersion objective with instrument minimized framerate.

### MGA array plasmid construction

Single copy malachite green aptamer-containing plasmids were derived from pcDNA3.1(+) (Thermo), digested with *BamHI* and *NotI* and similarly cut acGFP from pAcGFP-N1-SialT^(28)^ (pAcGFP1-N1-SialT was a gift from Lei Lu (Addgene plasmid #87324; http://n2t.net/addgene:87324; RRID:Addgene_87324) was inserted to create pcDNA3.1-acGFP. pcDNA3.1-acGPF was digested with *AgeI* and *XbaI* and inserted a similarly digested PCR product of mCDK6 from pcDNA3.1 mouse cdk6 wt (pcDNA3.1-mouse cdk6 wt was a gift from Martine Roussel (Addgene plasmid #75170; http://n2t.net/addgene:75170; RRID: Addgene_75170). Amplification was achieved using 5’-ATATATACCGGTACCATGGAGAAGGACAGCCT-3’ and 5’-ATATATTCTAGAATCAGGCTGTGTTCAGCTCC-3’, resulting in pcDNA3.1-mouse *Cdk6 wt*. The single copy malachite green aptamer^(18)^ was inserted by digesting pcDNA3.1acGFP with *NheI* and *BamHI* and inserting an annealed oligo pair, 5’– CTAGCGGATCCCGACTGGCGAGAGCCAGGTAACGAATGGATCC-3’ and 5’-GATCCGGATCCATTCGTTACCTGGCTCTCGCCAGTCGGGATCC-3’ with compatible overhangs. The resulting vector, pcDNA3.1-1xMGA-acGFP was digested with *AgeI* and *XbaI* and inserted a similarly digested PCR product of mCDK6 from pcDNA3.1 mouse cdk6 wt. Amplification was achieved using 5’-ATATATACCGGTACCATGGAGAAGGACAGCCT-3’ and 5’-ATATATTCTAGAATCAGGCTGTGTTCAGCTCC-3’, resulting in pcDNA3.1-1xMGA-mouse *Cdk6 wt*. pcDNA3.1-6xMGA-acGFP was made by digesting pcDNA3.1acGFP with *NheI* and *AflII* and inserting similarly cut 6xMGA^(29)^ PCR amplified from pUC57-6xMGA (Genscript) with 5’-ATATATGCTAGCTAGATGGTGTTTTGGTTTGG-3’ and 5’-ATATATCTTAAGCGAATTCGGATCCGCG-3’. pcDNA3.1-6xMGA-mouse Cdk6 was made by digesting pcDNA3.1-6xMGA-acGFP with *AgeI* and *XbaI* and inserted a similarly digested PCR product of mCDK6 from pcDNA3.1 mouse cdk6 wt. Amplification was achieved using 5’-ATATATACCGGTACCATGGAGAAGGACAGCCT-3’ and 5’-ATATATTCTAGAATCAGGCTGTGTTCAGCTCC-3’, resulting in pcDNA3.1-6xMGA-mouse *Cdk6*.

## Supporting information

Live cell tracking of cellular granules

granule dissolution with 1,6 Hexanediol

## ASSOCIATED CONTENT

**Supporting Information**.

## ACKNOWLEDGMENT

This work was supported by the National Institutes of Health (R01GM116991 to J.M.H.) and (R01MH109026 to GJB).

## TOC GRAPHIC

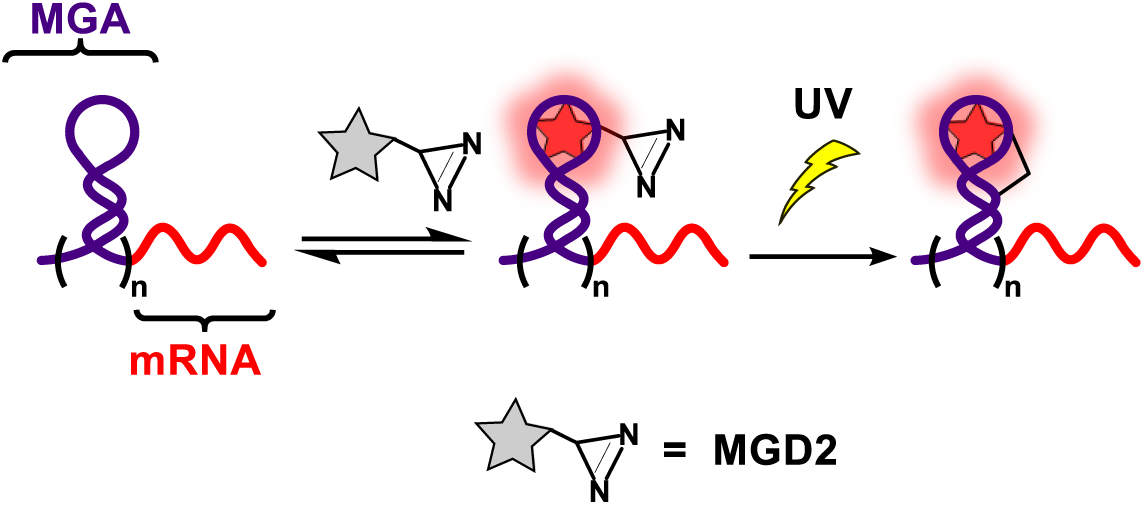

