## Supplemental Information for "Imaging and tracking mRNA in live mammalian cells via fluorogenic photoaffinity labeling"

**Table of Contents**

**Synthesis of leucomalachite green diazirine**	S2

**Synthesis of Malachite green diazirine (MGD2)**	S2

**Determination of MGD2 selectivity in cellular RNA**	S2-S3

**Tabular data fluorescence output of MGA array**	S3

**Tabular data for UV dependent fluorescence enhancement**	S3

**Tabular data for fluorescence of RNA foci**	S4

**Tabular data for Fluorescence signal comparison of GFP and MGA/MGD2**	S4

**Synthesis of MGD2**

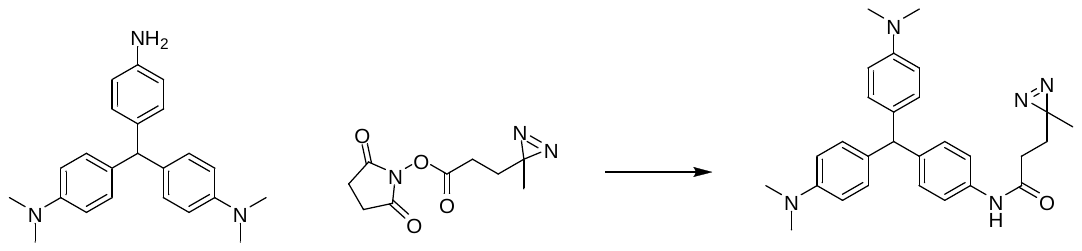

**Leucomalachite green diazirine**. The synthesis of *p-*amino-leucomalachite green was performed using the protocol described by Deng and coworkers.^1^ In an oven dried 5 mL flask, *p*-amino-leucomalachite green (25.0 mg, 72.7 µmol) was dissolved in 1 mL of dry pyridine under inert gas. To this solution, 1.2 eq. of NHS-diazirine (19.6 mg, 86.83 µmol) was added and the solution was stirred at room temperature overnight. The reaction was then concentrated under reduced pressure to obtain dark green oil. The crude oil was dissolved in minimal amount of methanol and loaded on a preparative TLC with 30% EtOAc/MeOH as a mobile phase. The product band was scraped off from the preparative TLC and the product was filtered from the silica using MeOH and dried under reduced pressure. Yield = 11.6 mg, 35.31%. ^1^H NMR (500 MHz, DMSO) δ 0.99 (s, 3H), 1.64 (t, 2H, *J* = 7.57 Hz), 2.20 (t, 2H, *J* = 7.3 Hz), 3.04 (s, 12H), 5.63 (s, 1H), 7.03 (d, 2H, *J* = 8.3 Hz), 7.24 (d, 4H, *J* = 7.3), 7.52-7.69 (m, 6H), 10.15 (s, 1H). ^13^C NMR (500 MHz, DMSO) δ 19.36, 25.76, 29.53, 30.69, 44.52, 53.93, 119.21, 129.06, 130.14, 137.68, 169.75. LRMS (ESI-TOP) *m/z* Calcd for C_28_H_33_N_5_O [M + H]^+^ 456.2758; Found 456.2746

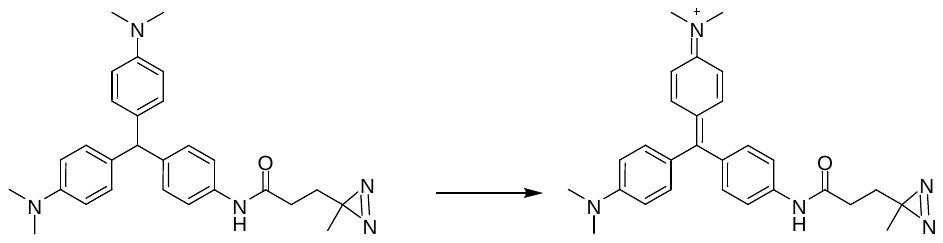

**Malachite green diazirine (MGD2)**. 11.64 grams (25.55 µmol) of leucomalachite green diazirine was dissolved in 20% MeOH/EtOAc. To this solution, 1.2 eq. of chloranil (7.54 mg, 30.66 µmol) was added and the solution was stirred at room temperature for 3 h. The dark green solution was concentrated under reduced pressure and flash column purified using EtOAc to remove excess chloranil. Then mobile phase was then switched to 50% MeOH/DCM to isolate the crude product. The collected crude product was concentrated under reduced pressure and further purified by preparative TLC using 20% MeOH/DCM as mobile phase. Yield = 10.30 mg, 88.68%. ^1^H NMR (500 MHz, DMSO) δ 1.03 ( s, 3H), 1.69 (t, 2H, *J* = 7.1 Hz), 2.37 (t, 2H, *J* = 7.3 Hz), 3.14 (s, 6H), 3.27 (s, 6H), 7.07 (d, 4H, *J* = 8.8 Hz), 7.31 (t, 6H, *J* = 9.3 Hz), 7.94 (d, 2H, *J* = 8.3 Hz). ^13^C NMR (500 MHz, DMSO) δ 19.35, 25.70, 28.93, 29.37, 40.39, 48.50, 113.63, 118.52, 126.24, 133.20, 136.25, 140.06, 156.23. LRMS (ESI-TOP) *m/z* Calcd. for C_28_H_32_N_5_O^+^ [M]^+^ 454.2601; Found 454.2595

**Determination of MGD2 selectivity in cellular RNA.** To determine the selectivity of MGD2 labeling, we spiked in 6MGA-mGFP (200 ng) into cellular RNA extracted from HeLa cells (400 ng) in 1x PBS. To this solution, MGD2 (50 µM final concentration) was added and the solution was incubated for 20 min at room temperature. After 10 min of UV irradiation, the sample was mixed with equal volume of 50% glycerol solution and heated to 70 ⁰C to denature the RNA. This sample was loaded on to 1% agarose gel containing 1% v/v Clorox® bleach^2^ and Sybr Gold (Thermofisher). After running the gel for 1 h at a constant 100 V, the gel was frozen with dry ice for 10 min and imaged using GE Amersham Typhoon (Supplementary Fig 1). This freezing step enhanced the fluorescence output of the malachite green molecule. Analysis of this gel indicates that the MGD2 selectively labeled the aptamer functionalized mRNA and did not have any detectable labeling of cellular RNA.

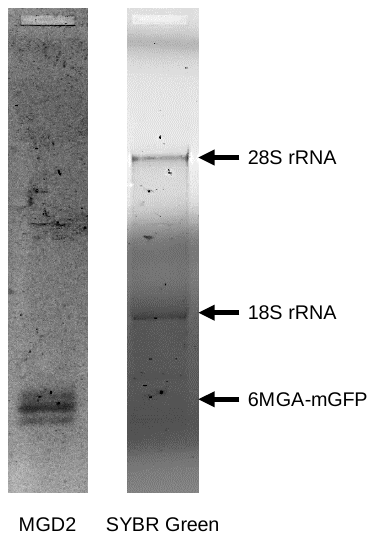

**Supplementary Fig 1.** Selective labeling of MGA functionalized mRNA in the presence of cellular RNA extracted from HeLa cells.

**Table S1.** Tabular data fluorescence output of MGA array

|  | MGD2 | 1xMGA-mGFP + MGD2 | 6xMGA-mGFP + MGD2 |
| --- | --- | --- | --- |
| mean | 6.25 | 1570 | 6454 |
| stdev | 0.96 | 73 | 150 |

**Table S2.** Tabular data for UV dependent fluorescence enhancement of 1xMGA-mRNA and control mRNA compared to MGD2 in 1xPBS

|  | 1xMGA-mGFP | | control mRNA | |
| --- | --- | --- | --- | --- |
| UV irradiation time (min) | mean | stdev | mean | stdev |
| 0 | 264 | 63 | 5.4 | 1.8 |
| 5 | 167 | 38 | 2.67 | 0.76 |
| 10 | 135 | 35 | 2.22 | 0.25 |
| 15 | 148 | 36 | 2.39 | 0.98 |

**Table S3.** Tabular data for fluorescence of RNA foci

|  | no transfection | Transfected with control RNA | 1x MGA | 6x MGA |
| --- | --- | --- | --- | --- |
| Lower quartile | -0.119 | 1.42 | 14.6 | 40.6 |
| Median | 0.771 | 3.28 | 26.9 | 50.3 |
| Upper quartile | 3.18 | 4.26 | 43.1 | 58.9 |
| 5% Percentile | -0.263 | -2.63 | 8.81 | 21.9 |
| 95% Percentile | 6.72 | 6.09 | 55.5 | 68.3 |
| Mean | 1.66 | 2.74 | 29.7 | 48.2 |

**Table S4** Tabular data for fluorescence signal comparison of GFP-tagged cellular protein and MGD2 labeled RNA granules

| Time (min) | Protein | | RNA | |
| --- | --- | --- | --- | --- |
|  | Mean | SEM | Mean | SEM |
| 0 | 0.955 | 0.0116 | 0.756 | 0.220 |
| 1.334 | 0.952 | 0.0106 | 0.783 | 0.154 |
| 2.668 | 0.985 | 0.00747 | 0.846 | 0.112 |
| 4.002 | 0.974 | 0.00603 | 0.895 | 0.0371 |
| 5.337 | 0.940 | 0.0187 | 0.939 | 0.0231 |
| 6.671 | 0.982 | 0.0114 | 0.950 | 0.0394 |
| 8.005 | 0.957 | 0.0197 | 0.939 | 0.0284 |
| 9.339 | 0.972 | 0.0187 | 0.959 | 0.0151 |
| 10.673 | 0.974 | 0.00936 | 0.924 | 0.0492 |
| 12.007 | 0.944 | 0.0119 | 0.955 | 0.0322 |
| 13.342 | 0.944 | 0.0175 | 0.935 | 0.0344 |
| 14.676 | 0.924 | 0.0112 | 0.928 | 0.0603 |
| 16.01 | 0.942 | 0.0073 | 0.934 | 0.0540 |
| 17.344 | 0.908 | 0.0284 | 0.980 | 0.00924 |
| 18.678 | 0.873 | 0.0361 | 0.980 | 0.0133 |
| 20.012 | 0.799 | 0.0577 | 0.959 | 0.0226 |
| 21.347 | 0.708 | 0.0556 | 0.960 | 0.0098 |
| 22.681 | 0.622 | 0.155 | 0.970 | 0.0116 |
| 24.015 | 0.509 | 0.122 | 0.897 | 0.0419 |
| 25.349 | 0.362 | 0.129 | 0.928 | 0.0411 |
| 26.683 | 0.330 | 0.0933 | 0.933 | 0.0347 |
| 28.017 | 0.259 | 0.0964 | 0.960 | 0.00347 |
| 29.352 | 0.199 | 0.0863 | 0.938 | 0.0314 |
| 30.686 | 0.188 | 0.0906 | 0.948 | 0.0164 |

1. Xing, W.; He, L.; Yang, H.; Sun, C.; Li, D.; Yang, X.; Li, Y.; Deng, A., Development of a sensitive and group‐specific polyclonal antibody‐based enzyme‐linked immunosorbent assay (ELISA) for detection of malachite green and leucomalachite green in water and fish samples. *Journal of the Science of Food and Agriculture* **2009,** *89* (13), 2165-2173.

2. Aranda, P. S.; LaJoie, D. M.; Jorcyk, C. L., Bleach gel: a simple agarose gel for analyzing RNA quality. *Electrophoresis* **2012,** *33* (2), 366-369.
